# A global assessment of how biodiversity enhances agroecosystem function

**DOI:** 10.64898/2026.09.15.751770

**Authors:** Olivia F. Morris, James M. Bullock, Emma Moffett, Guy Woodward, William D. Pearse

**Affiliations:** Georgina Mace Centre for the Living Planet, Department of Life Sciences, Imperial College London, Silwood Park Campus, Ascot, UK, SL5 7PY; UK Centre for Ecology & Hydrology, Wallingford, UK, OX10 8BB; Department of Geography, King’s College London, London, UK, WC2B 4BG; Alan Turing Institute, British Library, 96 Euston Road, London, UK, NW1 2DB

## Abstract

1. Agriculture is the most significant driver of global land-use change, placing increasing pressure on biodiversity. Balancing rising food demand with the need to conserve biodiversity is therefore an urgent challenge, especially as biodiversity can underpin key ecosystem functions that support agricultural systems. A greater general understanding of how biodiversity influences agroecosystem function is critical for developing agricultural practices that benefit both agricultural production and wider ecosystem processes.
2. Using a global dataset of 184 biodiversity-agroecosystem function relationships covering agricultural and silvicultural systems, we developed a classification system for different biodiversity-enhancing approaches, ranging from the planting of multiple crop species to creating tree islands or flower strips, providing the first comprehensive assessment of how these approaches affect agroecosystem function.
3. We found biodiversity enhancement generally had a positive, or at least neutral, association with multiple agroecosystem functions, indicating that it contributes to agricultural sustainability. Crop diversification had the greatest impact on key agroecosystem functions compared to other practices, likely due to the low heterogeneity already present in agricultural systems and the scale at which crop diversification is applied and measured. Overall, biodiversity’s largest effect was on multifunctionality, highlighting its role in supporting the simultaneous delivery of multiple agroecosystem functions.
4. Variability in relationships among systems and agroecosystem function categories suggests that no single biodiversity-enhancing approach will universally benefit all functions simultaneously. Instead, a combination of approaches, assessed through their contribution to overall multifunctionality, is likely to be most effective, ensuring that agricultural sustainability claims reflect benefits across systems.
5. Policies should incentivise biodiversity-enhancing management approaches as effective strategies to improve agroecosystem multifunctionality, benefiting both agricultural productivity and the wider ecosystem.

## Introduction

High biodiversity can enhance the functions, structures, and processes within ecosystems underpinning the services humans rely upon (Hooper *et al*., 2005; Tilman *et al*., 2014; van der Plas, 2019). However, biodiversity is in rapid decline globally, threatening the stability of ecosystems, their functioning, and the services they provide (IPBES, 2019; Pfenning-Butterworth *et al*., 2024). Addressing this decline is a pressing concern, with a growing awareness of the need to halt and reverse biodiversity loss (Leclère *et al*., 2020). One of the primary drivers of biodiversity loss is land-use change, particularly the rapid expansion and intensification of agriculture (Newbold, 2018; Beckmann *et al*., 2019; IPBES, 2019; García-Vega *et al*., 2024). Indeed, by 2050, food production is projected to need to increase by around 50% to sustain a growing global population (Alexandratos and Bruinsma, 2012; van Dijk *et al*., 2021). Achieving biodiversity targets within this context is challenging, as efforts to conserve nature often conflict with the growing demand for food (Ortiz *et al*., 2021). Balancing these competing demands is complex but will include decisions about whether and where land should be prioritised for food production, conserved for nature, or managed to support both objectives via some form of trade-off optimisation (Fischer *et al*., 2014; Woodcock *et al*., 2025). A key part of this challenge is therefore understanding how biodiversity influences ecosystem functioning within agricultural systems, where the primary focus is typically on maximising yield and production, yet long-term productivity and resilience also depend on the sustained delivery of these functions and services (Bullock *et al*., 2017).

Sustainable agricultural practices have gained increasing attention as a means to align food production with ecological sustainability (FAO, 2019). While this can take many forms, typically the focus is to prioritise the role of biodiversity in maintaining or increasing food production, while minimising environmental impacts and enhancing ecosystem service delivery (Thrupp, 2000; Scherr and McNeely, 2008; Muhie, 2022; Rehman *et al*., 2022; Sher *et al*., 2024). A growing body of evidence demonstrates positive links between biodiversity and ecosystem function within agricultural systems, both through above- and below-ground diversity (Altieri, 1999; Swift *et al*., 2004; Moonen and Bàrberi, 2008; Gonzalez *et al*., 2020; Tamburini *et al*., 2020; Woodcock *et al*., 2025). For example, plant biodiversity can influence soil microbial diversity, helping to regulate decomposition and nutrient cycling, which in turn supports crop yields (Thiele-Bruhn *et al*., 2012; Cappelli *et al*., 2022; Zhang *et al*., 2023). Similarly, features such as hedgerows and flower strips can support pollinators or encourage predators of parasites and pests either directly or through soil-mediated interactions (Tschumi *et al*., 2015; Alison *et al*., 2022; Kowalska *et al*., 2022). Additionally, practices such as rotational grazing can improve soil health and nutrient cycling by breaking down plant material and increasing nutrient availability, which in turn supports biodiversity in agroecosystems and contributes to ecosystem functioning and the provision of services (Teague and Kreuter, 2020; Sher *et al*., 2024; Stanley *et al*., 2024).

Despite growing recognition of the potential for biodiversity-led practices to support more sustainable agricultural systems, their implementation remains limited (Duru *et al*., 2015). Food systems frequently prioritise short-term gains in production over long-term ecosystem health (EEA, 2022; Thierfelder and Mhlanga, 2022; Brooker *et al*., 2023). This is in part likely due to the benefits of biodiversity for production varying across geographical and agricultural contexts, making it challenging to develop universal guidelines (Bommarco *et al*., 2013; Wittwer *et al*., 2017; Wan *et al*., 2018; Ji *et al*., 2023; Jones *et al*., 2023). It is evident that the impact of biodiversity on agroecosystem functions is influenced by crop type (dos Santos *et al*., 2021), climate (Ortiz *et al*., 2021; Jones *et al*., 2023), and the spatial (Brooker *et al*., 2023) and temporal scale of an intervention or study (Buzhdygan and Petermann, 2023). However, these uncertainties highlight the need to assess the impact of biodiversity on agroecosystem functions across a broad range of systems and contexts in order to disentangle patterns.

Here, we use a global dataset comprising over 13,100 measurements from 184 biodiversity-agroecosystem function relationships within agricultural and silvicultural systems. We compare a range of biodiversity-enhancing management approaches considered sustainable, from multiple crops to rewilding strategies and reduced grazing intensity, as well as a number considered unsustainable, such as fertiliser use. We define agroecosystem functions as ecosystem processes that take place within agricultural ecosystems; however, we recognise that many of these functions also underpin, or may be described as, ecosystem services. By categorising these approaches and testing the consistent strength and direction of biodiversity’s impact on agroecosystem functions across management practices, we aim to identify general patterns that can inform the development of more sustainable and biodiversity-led agricultural systems. Although biodiversity enhancement is generally assumed to improve agroecosystem function, the evidence to date is inconsistent, and it remains unclear when and why such relationships are observed or not. We seek to resolve this uncertainty by identifying the contexts, management strategies, and agroecosystem function categories in which biodiversity enhancement most strongly supports agricultural sustainability.

## Materials and methods

Using collated data covering cropland, pastureland, and forestry systems, we examined how various biodiversity-enhancing approaches affect agroecosystem functions, such as food and feed production, water quality, or the regulation of pests. Practices were classified as either biodiversity-led sustainability approaches, where biodiversity enhancement is an explicit management goal (e.g., enhanced hedgerows, multiple crop species), or as comparison practices, encompassing both unsustainable management (e.g., fertiliser use) and incidental biodiversity change, where biodiversity variation occurs indirectly rather than through deliberate management. After scaling the data to make it comparable across different systems and contexts, we fit a Bayesian hierarchical model, accounting for the uncertainty within and across studies. All analyses were conducted in R (version 4.4.0; *R Core Team,* 2024).

### Data collection

We extracted data from a dataset compiled by Moffett et al. (in press), which was derived from a stratified systematic review of the effects of biodiversity on ecosystem functions and services, along with associated metadata, such as study design and methodological details. Their review used the Web of Science to identify studies (published up to 2024) and resulted in a dataset of 545 studies. For our analysis, we filtered this dataset to include only studies conducted in agricultural settings, specifically within cropland, pastureland, and plantations. This resulted in a final subset of 28 studies included in our analysis. While limited, they collectively contribute 184 separately measured relationships since individual studies frequently measured multiple relationships, each of which had at least six data points. We grouped agroecosystem functions within the dataset according to the classification in Moffett et al. (in press), which follows the IPBES framework for ecosystem services (e.g., food and feed provision, IPBES, 2019), while also distinguishing key ecosystem functions. We also make use of this classification as it provides a standardised approach that enables cross-study comparisons.

We developed a classification system to categorise each study based on the specific sustainable agriculture practice implemented to enhance biodiversity, defining sustainability in terms of its impact on biodiversity (hereafter referred to as ‘sustainable practices’). Although sustainable agriculture can take many forms, we define it here as biodiversity-led practices that prioritise biodiversity within an agricultural system (Muhie, 2022). We also recognise that this definition overlaps with related terms such as regenerative agriculture or ecological intensification; however, these share similar goals of improving ecosystem function and service provision (Newton *et al*., 2020; Giller *et al*., 2021; Sher *et al*., 2024). Our classification system builds upon existing classification frameworks (see LaCanne and Lundgren, 2018; Tamburini *et al*., 2020; López Rodríguez *et al*., 2024; Storkey *et al*., 2024) but extends them to capture the full range of biodiversity-driven interventions within agricultural systems from our dataset. This enables a standardised comparison of how different management strategies influence agroecosystem function across different agricultural contexts, to test whether biodiversity-led strategies vary in their effectiveness. We therefore included the following sustainable practices:

1. ‘*Non-crop diversification’*, where biodiversity is increased through practices such as planting flower strips or maintaining wild habitats;
2. ‘*Crop diversification’*, which included approaches such as intercropping and polyculture to enhance biodiversity within the cultivated system;
3. ‘*Combined diversification’*, where both crop and non-crop diversification strategies were applied in unison;
4. ‘*Organic practices*’, characterised by the reduced or non-use of agrochemical pesticides and fertilisers; and
5. ‘*Grazing management’*, where livestock production intensity was reduced to promote biodiversity. Finally, we included a broader grouping (‘*other’*) of categories for studies where biodiversity change was measured but not explicitly driven by sustainable management interventions. These included:
6. *climatic gradients*;
7. *land-use types*;
8. *intensive grazing*, intensive agricultural systems where biodiversity was measured but not intentionally enhanced;
9. *fertiliser* application, where conventional (non-sustainable) inputs were used.

These studies do not typically align with biodiversity-led sustainability practices, with some observational rather than management-led, but were retained to compare effects. As we highlight below, some of these categories are better-studied than others; assessing the size of the evidence-base for each kind of practice was one of our study aims and helps to guide future work. We emphasise that our use of a Bayesian hierarchical modelling approach (see below) means that our findings are robust to differences in sampling; the estimates of practices with less data simply have more uncertain estimates.

Lastly, we accounted for temperature as a key environmental variable. Using the ERA5 climate reanalysis dataset (Hersbach *et al*., 2020), we extracted the mean near-surface (2m) air temperature for the GPS coordinates corresponding to each study for the year (or years) in which each study was conducted, to capture potential environmental variation across studies.

### Analysis

To estimate general linear trends and whether changes in biodiversity are associated with an increase or decrease in agroecosystem function, we used a Bayesian hierarchical model implemented in *rstanarm* (Goodrich *et al*., 2025). The model captures biodiversity effects (slopes), which represent the strength and direction of biodiversity’s impact on agroecosystem function. We modelled a linear relationship, due to typically low biodiversity in agricultural systems and where increases in biodiversity are unlikely to reach levels where agroecosystem functions saturate (Weisser *et al*., 2017). To examine general patterns, while investigating the role of key variables, we modelled the interaction between biodiversity change with hierarchical terms included additively, reflecting the nested structure of the data where multiple measurements occur within studies and shared contexts. These included the above sustainability classifications, agroecosystem function categories, farming type (crop, livestock, forestry), study duration (short-term: 0–1 years, medium-term: 1–5 years, long-term: 5–10 years, and very long-term: over 10 years), mean temperature of the study, and the spatial extent of each study (standardised to square metres and log-transformed to account for large differences across studies). We also included a hierarchical term for each unique biodiversity-agroecosystem function relationship, allowing us to capture across- and within-study slope variation. To enable clearer interpretation of model estimates, we set the reference categories (the groups to which all other estimates are compared) as the category with the lowest biodiversity effect (i.e., the shallowest slope): *land-use types (other)* for the agricultural practices and *terrestrial carbon sequestration* for the agroecosystem function metric. Thus, categories that are significant are significantly *greater* in their impacts than either of these references. To ensure comparability across studies and account for heterogeneity in the measurement units of biodiversity and agroecosystem function values, all continuous terms were standardised. They were scaled within each biodiversity-agroecosystem function relationship by centring on the mean and dividing by two standard deviations, following Gelman (2008). This approach enables comparisons between continuous and categorical predictors, allowing for more interpretable estimates of relative effect sizes. We also do not report intercept estimations from the model, as scaling centres the data at zero and limits interpretability.

## Results

Data from 28 studies were analysed, resulting in 184 distinct measures of how agroecosystem function varies in response to biodiversity change across agricultural systems globally (Fig. 1). The majority of studies incorporated sustainability measures (n=23), while 8 included no explicitly biodiversity-led sustainable practices (i.e., the *other* category). Three studies contained both sustainable and non-sustainable observations. *Non-crop diversification* was the most commonly applied practice, appearing in 11 studies and accounting for 68 biodiversity-agroecosystem function relationships (Fig. 2). The distribution of the data across sustainability practices also highlights differences in representation (Fig. 3), with *non-crop diversification* also encompassing the largest amount of associated data points (n=3014).

**Figure 1.**
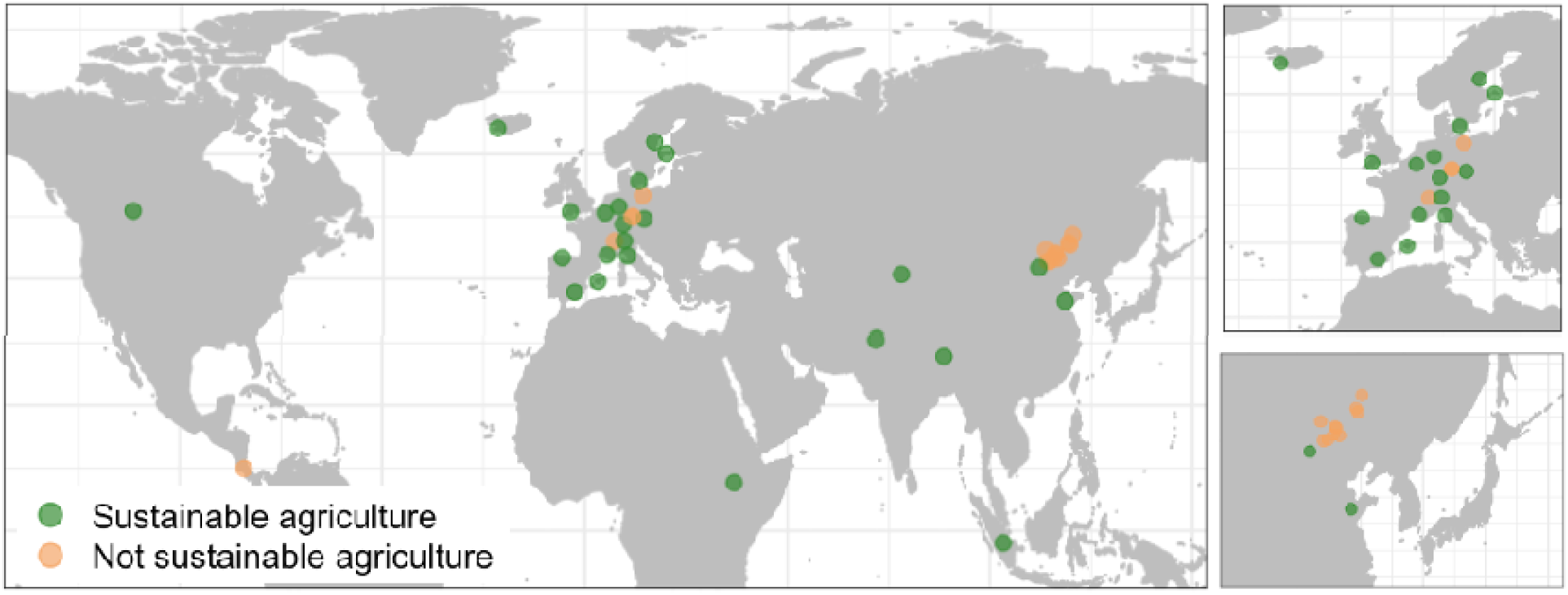
Global distribution of study locations within the dataset. The global distribution of the data coloured by the sustainable biodiversity-led practice applied to measure biodiversity change on agroecosystem function. Sustainable practices are shown in green, with other practices not related to sustainable agriculture, but where biodiversity varied within an agricultural system, represented in orange. The data are biased towards the Northern Hemisphere, as well as regions such as Europe and China, as shown by the magnified sections.

**Figure 2.**
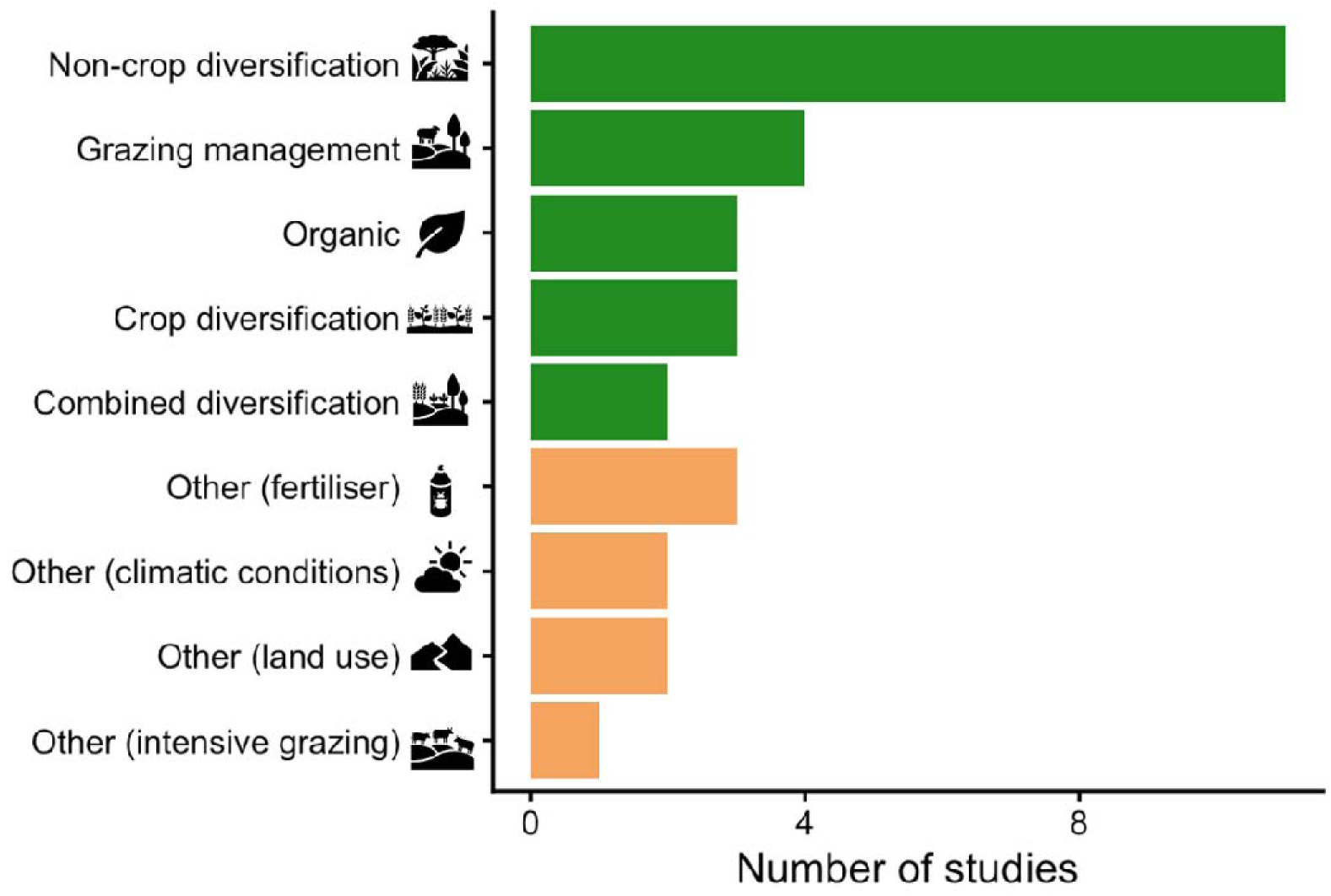
Non-crop diversification was the most studied practice across biodiversity-enhancing agricultural practices. The number of studies within our dataset measuring biodiversity and agroecosystem function in agricultural systems, along with the types of manipulations performed. *Sustainable* biodiversity practices are shown in green, while *other* practices are depicted in orange. The most frequently tested practice was *non-crop diversification*, a sustainable practice that involved enhancing biodiversity through practices such as tree islands, flower strips, or hedgerows.

**Figure 3.**
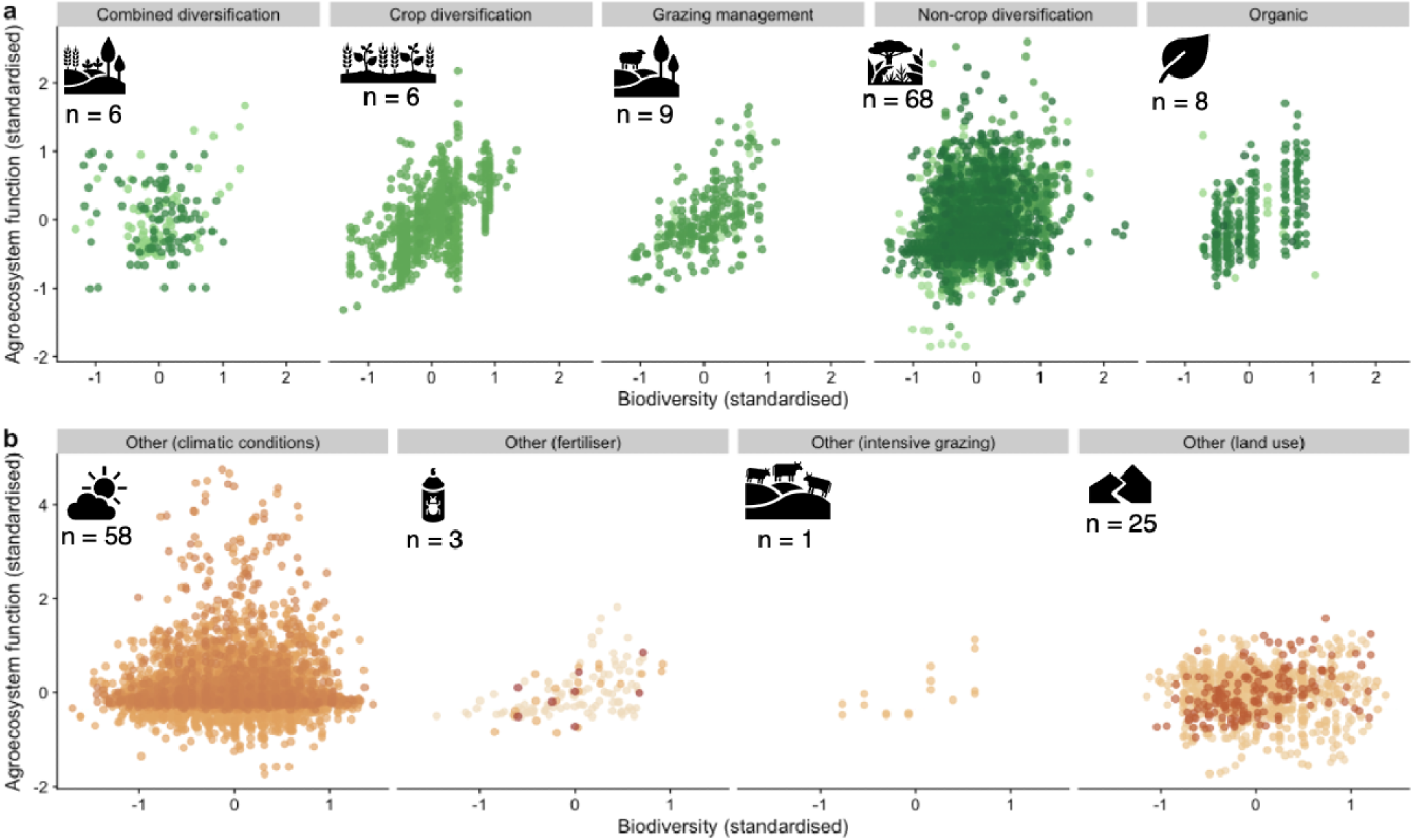
The relationship between standardised biodiversity and agroecosystem function across agricultural practices. The standardised data points showing the relationship between biodiversity change (x-axis) and agroecosystem function (y-axis) for each practice. Panel (a) shows the data for *sustainable* practices (shades of green), while panel (b) shows the data for *other* agricultural practices where manipulated biodiversity was measured (shades of orange). The number (n) of individual biodiversity-agroecosystem function relationships is given under each category icon, with the data points shown in different shades to distinguish individual relationships. These data highlight the variation in the relationships across sustainability practices, with *non-crop diversification* having the most associated data.

### Biodiversity effects within agricultural practices

Among the agricultural practices, *crop diversification* exhibited the greatest positive biodiversity effect, followed by climatic differences (*other (climatic conditions*)) and fertiliser management (*other (fertiliser)*; Fig. 4a). Sustainable practices, such as *combined diversification, grazing management,* and *non-crop diversification*, were associated with weak or negligible biodiversity effects. Other predictors within the model generally showed minimal influence on the slope (but we emphasise that many were under-represented and so perhaps under-studied). Notable exceptions were livestock systems, where the biodiversity effect was more positive relative to crops and forestry (0.71,80% CrI:0.39 - 1.04), and long-term studies lasting between 5 and 10 years, where the biodiversity effect was also more positive (0.57, 80% CrI: 0.11 - 1.03) compared to short-term studies (0-1 years). However, very long-term studies (greater than 10 years) were associated with a more negative effect relative to short term studies ( - ), suggesting a non-linear relationship over time. The full model, including all predictors, is provided in Supplementary Material Table 1.

**Figure 4.**
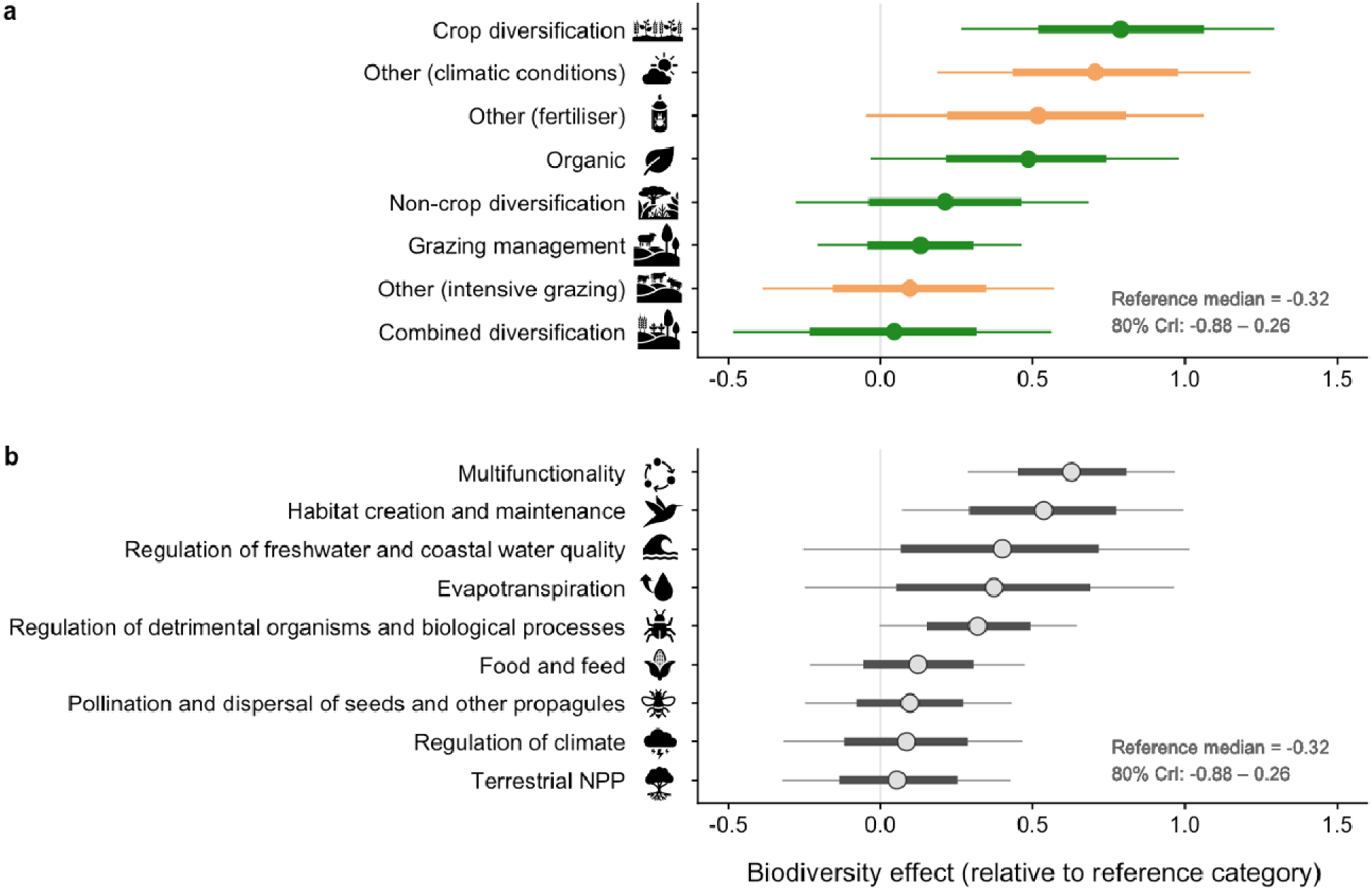
Crop diversification and multifunctionality are associated with the largest positive biodiversity effects. Model estimates for the biodiversity effects from posterior distributions, showing the median with 50% (bold line) and 80% (light line) credible intervals. Panel (a) shows the effects for agricultural practices, and panel (b) shows the effects for agroecosystem function categories. Values represent the change in biodiversity effect relative to the reference categories: *terrestrial carbon sequestration*, *other (land use)*, *crop systems*, and *short-term (0–1 year)* study length, which had biodiversity effects statistically indistinguishable from zero. Each panel shows the reference median with an 80% credible interval that crosses zero, indicating no effect (see Supplementary Material Table 1 for full model coefficients). For example, conditional on other moderators being at their reference levels, the estimated biodiversity effect for multifunctionality is −0.32+0.63 = 0.31, where −0.32 is the reference biodiversity effect (*terrestrial carbon sequestration;* not credibly different from zero) and 0.63 is the additional estimated effect of *multifunctionality*. *Crop diversification* had the greatest biodiversity effect among agricultural practices, and among agroecosystem function categories, *multifunctionality* (where studies measured more than one agroecosystem function simultaneously) had the largest biodiversity effect.

### Variation across agroecosystem function categories

Across agroecosystem function categories, *multifunctionality,* where studies measured more than one function simultaneously, showed the strongest positive biodiversity effect (Fig. 4b). This was followed by *habitat creation and maintenanc*e, which also had a significant positive biodiversity effect. In contrast, *terrestrial carbon sequestration* (the reference category), *terrestrial NPP* and *regulation of climate* showed the lowest biodiversity effects; however, these were not credibly different from zero.

## Discussion

Although the importance of biodiversity in supporting ecosystem functions is well acknowledged (Swift *et al*., 2004; Haines-Young and Potschin, 2010; Brockerhoff *et al*., 2017; IPBES, 2019; Weiskopf *et al*., 2022), it is still unclear how broadly biodiversity affects agroecosystem function across systems, which makes it challenging to develop targeted strategies for improving agricultural sustainability (Brooker *et al*., 2021; Ortiz *et al*., 2021). Within this study, we explored the effects of biodiversity-enhancing agricultural practices on agroecosystem function. Consistent with previous research, our findings highlight the complexity of biodiversity-agroecosystem function relationships (Loreau *et al*., 2001; Bullock *et al*., 2021; Jones *et al*., 2023). In general, however, we find that biodiversity tends to have a positive or, at worst, neutral association across agroecosystem function. These findings reinforce that enhancing biodiversity can contribute to, and is unlikely to compromise, agricultural systems, improving functions that are both directly and indirectly beneficial to agricultural productivity. Non-significant relationships, however, do not indicate that agroecosystems are not providing these functions, but rather that their provision does not consistently increase with biodiversity gradients within these systems.

### Biodiversity effects across agricultural practices

Among the agricultural practices, crop diversification (i.e. where biodiversity is enhanced by incorporating multiple crop species through methods such as mixed cropping) had the greatest estimated positive biodiversity effect. Crop diversification data were represented by studies of food and feed production, biomass turnover rate, and terrestrial net primary production (NPP). This is consistent with previous studies showing that greater crop species diversity enhances soil quality, thereby improving yields and overall ecosystem productivity (Altieri, 1999; Moss *et al*., 2019; Tamburini *et al*., 2020; Mudare *et al*., 2025). Crop diversification may also have more pronounced effects on functions such as biomass turnover, as diverse cropping systems promote varied root structures and increase microbial activity in the soil, which accelerates organic matter decomposition and nutrient cycling (Martínez-García *et al*., 2018; Zhang *et al*., 2021; Williams *et al*., 2023). Such practices can therefore enhance yields both directly, through soil processes, but also indirectly, for example, by suppressing pests and reducing disease pressure (Mihrete and Mihretu, 2025), creating cascading benefits across the agroecosystem.

Notably, biodiversity was also estimated to be positively associated with agroecosystem functioning within fertilised systems. Although fertiliser use can have environmental costs and is often associated with intensive agricultural systems, it may provide short-term or context-specific benefits, particularly in nutrient-poor conditions (Andrey *et al*., 2014). This indicates that within fertilised systems, even small increases in biodiversity will yield larger improvements in agroecosystem function than solely relying on fertiliser inputs. Yet, while fertiliser use can yield immediate benefits, long-term effects can be detrimental, as continuous fertiliser use can lead to soil degradation, nutrient imbalances, and disruptions to soil microbial communities, ultimately impairing agroecosystem functioning (Kopittke *et al*., 2019; Asadu *et al*., 2024). This highlights a broader challenge for agricultural systems, in that practices that maximise short-term productivity may undermine long-term ecosystem health. Our results suggest that enhancing biodiversity within such systems may improve short-term agroecosystem functioning and potentially help mitigate some of these longer-term impacts; however, fertiliser use remains underrepresented in our dataset, limiting our ability to fully resolve these patterns. As a result, coordinated experiments that explicitly contrast short-term responses with longer-term outcomes (soil health, nutrient balance, pest dynamics), along with economic valuations, would be particularly valuable.

Surprisingly, many practices considered sustainable showed weak or negligible biodiversity effects, with additional increases in biodiversity yielding little further improvement in agroecosystem function. This may suggest that additional increases in biodiversity within these systems do not translate into further gains in agroecosystem function, reflecting a saturation point of provision. However, it may arise due to a potential mismatch in the spatial scale between the applied practices and the agroecosystem function measured, as local versus landscape-level processes can moderate the strength and detectability of biodiversity effects (Tscharntke *et al*., 2012; Duru *et al*., 2015; Brooker *et al*., 2023). For example, non-crop diversification practices, such as tree islands or enhancing hedgerows, are the most studied, yet are typically implemented at the field margin level, while agroecosystem functions are typically assessed at the broader field scale. In contrast, crop diversification is generally applied at the field scale, which may better align with the scale at which agroecosystem function is measured (Buzhdygan and Petermann, 2023; Arndt and Helming, 2025). Recent work shows that in-field diversification, such as introducing strips within crop fields, can lead to clear positive effects on biodiversity and associated agroecosystem function (Woodcock *et al*., 2025), whereas margin-based flower diversification benefits may decline sharply with distance from the field edge (Woodcock *et al*., 2016). These potential scale mismatches may therefore mask the benefits of sustainable practices. Beyond scale, biodiversity-agroecosystem function relationships may be further influenced by adjacent land use in ways that are difficult to fully account for, for example more intensive neighbouring fields (Williams *et al*., 2024). In particular, agroecosystem functions such as pollination and seed dispersal are strongly landscape-mediated (Kraus *et al*., 2025), suggesting that field-level analyses alone may systematically underestimate the broader value of biodiversity enhancement in agricultural landscapes, and highlighting the importance of accounting for spatial context in both management and measurement in these systems.

### Biodiversity effects across agroecosystem function categories

Across practices, biodiversity showed its largest positive effect on multifunctionality, suggesting that it supports the simultaneous delivery of multiple agroecosystem functions. This is also consistent with diverse communities providing functional complementarity or redundancy, which may buffer agroecosystems against trade-offs among functions rather than acting as a direct driver of any one process (Isbell *et al.,* 2011; Gimona *et al*., 2023). However, the specific functions included in measures of multifunctionality will differ among studies, which points to biodiversity’s integrative role rather than evidence for a particular mechanism. Biodiversity was also positively associated with improved habitat creation and maintenance, in that diverse communities may generate and stabilise the habitat structure they in turn depend on, which is particularly relevant given that increasing agricultural intensity tends to reduce habitat diversity (Lomba *et al*., 2022).

Effects on other agroecosystem functions were generally small, likely reflecting variation or processes not fully captured in our analyses. For example, climate regulation is largely driven by broader abiotic factors, and agricultural systems are already heavily modified in ways that could overshadow potential buffering effects (Brockerhoff *et al*., 2017; Saco *et al*., 2021; Stewart *et al*., 2022; Weiskopf *et al*., 2022). Small effects may also reflect how biodiversity itself was measured. In our dataset, the majority of biodiversity metrics were taxonomic, with relatively few functional and no phylogenetic measures (see Supplementary Figure 2), despite growing evidence that taxonomic diversity measures alone may not best reflect the functioning of ecosystems (Wood *et al*., 2015; Brooker *et al*., 2023). Biodiversity metrics incorporating functional traits or phylogenetic relationships have been shown to be reliable predictors of agroecosystem function (Cadotte *et al*., 2009; Tucker *et al*., 2019; dos Santos *et al*., 2021). As a result, expanding the use of these measures within agricultural studies could offer a greater understanding of how biodiversity drives different agroecosystem functions and helps address the current evidence gap on biodiversity responses to agricultural interventions (Bonfanti *et al*., 2025).

While factors such as spatial extent and environmental conditions can influence biodiversity-agroecosystem function relationships (Hooper *et al*., 2005; Moss *et al*., 2019; Gonzalez *et al*., 2020), these factors showed no consistent effect in our analysis, suggesting that the broad patterns we observed are relatively robust across study designs. Study duration, however, showed a non-linear pattern: relative to short-term studies the biodiversity–agroecosystem function relationship was more positive in studies lasting 5–10 years, but more negative in studies lasting more than 10 years. This may indicate that biodiversity effects change over long timescales, potentially strengthening over intermediate timescales before weakening or becoming more context dependent. We found, however, an uneven representation of agroecosystem function categories across agricultural types, management practices, and scales, which may confound observed patterns (see Supplementary Figure 1). An imbalance in the underlying data is also consistent with previous reviews of the field (Brooker *et al*., 2023; Buzhdygan and Petermann, 2023; Bonfanti *et al*., 2025), and our analysis further reveals vital gaps in the role of biodiversity within farming systems. Expanding coverage across agroecosystem functions, practices and the type of agriculture will be important for disentangling variation and strengthening general inferences regarding the role of biodiversity.

## Conclusions

Overall, our results highlight broader challenges in assessing the sustainability of agricultural systems, particularly as some practices are marketed as sustainable or regenerative despite limited evidence of consistent biodiversity and agroecosystem function benefits (Newton *et al*., 2020; Giller *et al*., 2021; Khangura *et al*., 2023). Without clear criteria and ground-truthing with evidence, such claims risk promoting practices that may not reliably enhance biodiversity or agroecosystem function in the long term. Across practices, we found that crop diversification had the strongest and most consistent positive biodiversity effects on agroecosystem function. In contrast, other practices showed weaker or more variable outcomes, likely reflecting differences in scale, management intensity, and the limited variability within already simplified agricultural systems. Notably, biodiversity’s largest effect across practices was on multifunctionality rather than a single function, indicating that it enables the simultaneous delivery of multiple agroecosystem functions within a system. These findings suggest that no single agricultural practice will universally enhance all agroecosystem functions. Instead, a combination of approaches, valued by their contribution to overall multifunctionality rather than gains in any one function, is more likely to be beneficial. Smaller-scale experimental studies already show that such combinations can create synergies among agroecosystem functions (Bullock *et al*., 2021), and tailoring these to context is therefore likely to be important for supporting agricultural sustainability. We therefore call for more targeted research into the effectiveness of both individual and combined biodiversity-enhancing practices on agroecosystem function. Clearer definitions of what constitutes sustainability, paired with robust evidence, will be essential for aligning agricultural management with global biodiversity goals and optimising their trade-offs.

## Notes

### Competing Interest Statement

The authors have declared no competing interest.

