## Supplementary Material for "A global assessment of how biodiversity enhances agroecosystem function"

### Supporting Information

**Supporting Information Table 1. Extended summaries of model coefficients and variances.** The full Bayesian hierarchical model terms, along with the random effect and residual variances. Values represent medians, 80% credible intervals (CrI), and standard deviations (SD). Interactions with standardised biodiversity (Biodiversity_value_x_scaled) relate to the biodiversity effect (slope).

| **Parameter** | **Median** | **10% CrI** | **90%** **CrI** | **SD** |
| --- | --- | --- | --- | --- |
| (Intercept) | 0 | -0.01 | 0.01 | 0 |
| Biodiversity_value_x_scaled | -0.32 | -0.88 | 0.26 | 0.45 |
| Biodiversity_value_x_scaled:Ecosystem_function_metricEvapotranspiration | 0.37 | -0.25 | 0.96 | 0.48 |
| Biodiversity_value_x_scaled:Ecosystem_function_metricFood_and_feed | 0.12 | -0.23 | 0.47 | 0.28 |
| Biodiversity_value_x_scaled:Ecosystem_function_metricHabitat_creation_and_maintenance | 0.54 | 0.07 | 1 | 0.36 |
| Biodiversity_value_x_scaled:Ecosystem_function_metricMultifunctionality | 0.63 | 0.29 | 0.97 | 0.27 |
| Biodiversity_value_x_scaled:Ecosystem_function_metricPollination_and_dispersal_of_seeds_and_other_propagules | 0.1 | -0.25 | 0.43 | 0.27 |
| Biodiversity_value_x_scaled:Ecosystem_function_metricRegulation_of_climate | 0.09 | -0.32 | 0.47 | 0.31 |
| Biodiversity_value_x_scaled:Ecosystem_function_metricRegulation_of_detrimental_organisms_and_biological_processes | 0.32 | 0 | 0.64 | 0.26 |
| Biodiversity_value_x_scaled:Ecosystem_function_metricRegulation_of_freshwater_and_coastal_water_quality | 0.4 | -0.26 | 1.01 | 0.49 |
| Biodiversity_value_x_scaled:Ecosystem_function_metricTerrestrial_NPP | 0.05 | -0.32 | 0.43 | 0.29 |
| Biodiversity_value_x_scaled:Ag_classificationCrop and non crop diversification | 0.04 | -0.48 | 0.56 | 0.41 |
| Biodiversity_value_x_scaled:Ag_classificationCrop diversification | 0.79 | 0.26 | 1.29 | 0.4 |
| Biodiversity_value_x_scaled:Ag_classificationGrazing management | 0.13 | -0.21 | 0.46 | 0.26 |
| Biodiversity_value_x_scaled:Ag_classificationNon crop diversification | 0.21 | -0.28 | 0.68 | 0.37 |
| Biodiversity_value_x_scaled:Ag_classificationOrganic | 0.49 | -0.03 | 0.98 | 0.39 |
| Biodiversity_value_x_scaled:Ag_classificationOther (climatic conditions) | 0.7 | 0.18 | 1.22 | 0.4 |
| Biodiversity_value_x_scaled:Ag_classificationOther (fertiliser) | 0.52 | -0.05 | 1.06 | 0.43 |
| Biodiversity_value_x_scaled:Ag_classificationOther (grazing intensity) | 0.1 | -0.39 | 0.57 | 0.37 |
| Biodiversity_value_x_scaled:temp_mean_scaled | 0.07 | -0.02 | 0.16 | 0.07 |
| Biodiversity_value_x_scaled:Ag_typeForestry | -0.01 | -0.11 | 0.09 | 0.08 |
| Biodiversity_value_x_scaled:Ag_typeLivestock | 0.71 | 0.39 | 1.04 | 0.25 |
| Biodiversity_value_x_scaled:Length_of_study1-5y | -0.12 | -0.25 | 0.01 | 0.1 |
| Biodiversity_value_x_scaled:Length_of_study10+ | -0.42 | -0.68 | -0.16 | 0.2 |
| Biodiversity_value_x_scaled:Length_of_study5-10y | 0.57 | 0.11 | 1.03 | 0.36 |
| Biodiversity_value_x_scaled:spatial_extent_m2_scaled | 0.01 | -0.09 | 0.11 | 0.08 |
| Random effect variance | 0.043 | 0.033 | 0.054 | 0.008 |
| Residual variance ($\sigma^{2}$) | 0.226 | 0.223 | 0.230 | 0.003 |

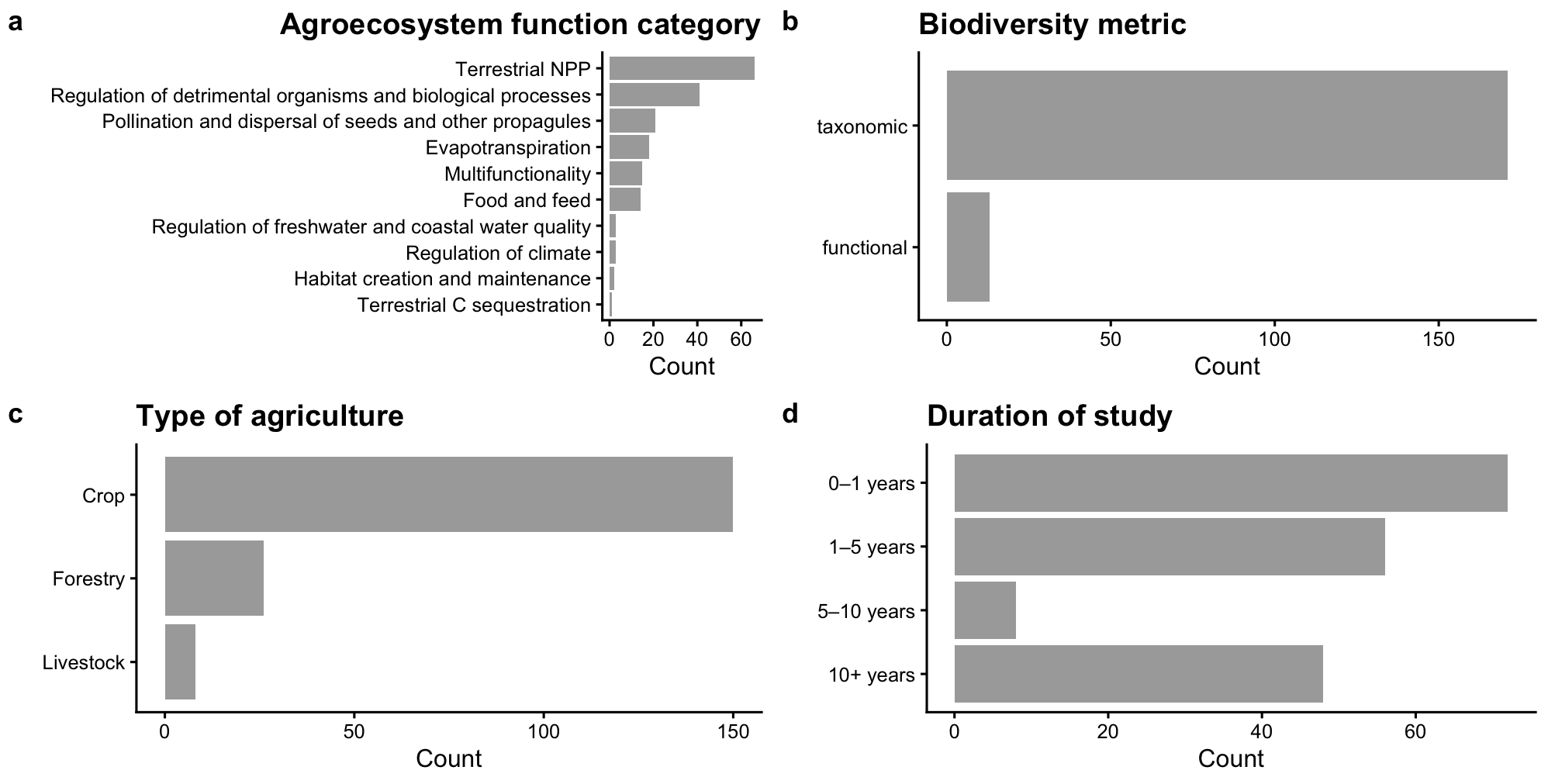

**Supporting Information Figure 1. Amount of data associated with the categorical variables in the dataset.** Each panel shows the frequency of observations (count: number of biodiversity-agroecosystem function relationships) within the data across each categorical variable in the model: (a) agroecosystem function category, (b) biodiversity metric type, (c) agroecosystem type, and (d) study duration.
